# Genome Compartmentalization with Nuclear Landmarks: Random yet Precise

**DOI:** 10.1101/2021.11.12.468401

**Authors:** Kartik Kamat, Yifeng Qi, Yuchuan Wang, Jian Ma, Bin Zhang

## Abstract

The three-dimensional (3D) organization of eukaryotic genomes plays an important role in genome function. While significant progress has been made in deciphering the folding mechanisms of individual chromosomes, the principles of the dynamic large-scale spatial arrangement of all chromosomes inside the nucleus are poorly understood. We use polymer simulations to model the diploid human genome compartmentalization relative to nuclear bodies such as nuclear lamina, nucleoli, and speckles. We show that a self-organization process based on a co-phase separation between chromosomes and nuclear bodies can capture various features of genome organization, including the formation of chromosome territories, phase separation of *A/B* compartments, and the liquid property of nuclear bodies. The simulated 3D structures quantitatively reproduce both sequencing-based genomic mapping and imaging assays that probe chromatin interaction with nuclear bodies. Importantly, our model captures the heterogeneous distribution of chromosome positioning across cells, while simultaneously producing well-defined distances between active chromatin and nuclear speckles. Such heterogeneity and preciseness of genome organization can coexist due to the non-specificity of phase separation and the slow chromosome dynamics. Together, our work reveals that the co-phase separation provides a robust mechanism for encoding functionally important 3D contacts without requiring thermodynamic equilibration that can be difficult to achieve.

## Introduction

Growing evidence has demonstrated that the three-dimensional (3D) organization of eukaryotic genomes plays essential roles in DNA-templated processes.^1–10^ Specifically, advancements in high-throughput sequencing and microscopic imaging have revealed sub-megabase, fine-scale structural features within individual chromosomes, including chromatin loops^11^ and topologically associating domains (TADs). ^12,13^ These structures can facilitate interactions between regulatory elements that are far apart in the genome to control gene expression. Significant progress has also been made in underpinning the molecular mechanisms that give rise to such structures. ^14–18^

However, at a global level, the presence of robust features of large-scale genome organization across chromosomes is debatable.^19–21^ General trends do exist, and different chromosomes tend to occupy preferred nuclear locations, with active and inactive chromatin generally residing near the nuclear interior and periphery, respectively.^22^ In fact, the relative distances of chromosomes to nuclear speckles are so precise that a slight variation on the order of 100 nm can lead to dramatic changes in gene expression levels.^23^ On the other hand, global genome organization also appears random. Variations can be readily seen in microscopic images that directly probe the spatial locations of chromosomes in individual cells.^24–26^ Chromosome radial positions are also not strictly inherited across cell cycles, ^27^ arguing against a significant functional role that might justify their maintenance.

A coherent mechanism that reconciles the various observations of global genome organization is currently missing. Phase separation has been proposed to drive the genomewide compartmentalization of euchromatin and heterochromatin.^28–31^ However, a block co-polymer model with attractive interactions between compartments of similar types fails to position heterochromatin towards nuclear envelope.^32,33^ In addition, a phase-separated liquid state is homogeneous and highly dynamic. It can hardly produce well-defined distances to explain the precise nature of genome organization. Several recent experimental techniques, including DamID,^34^ SPRITE^35^ and TSA-Seq, ^36^ have revealed close contacts between chro-mosomes and nuclear landmarks. These contacts are mediated by specific proteins,^37,38^ and could significantly impact the nuclear localization of chromatin and the radial positioning of chromosomes. They needed to be explicitly accounted for a complete mechanistic understanding of genome organization. ^39–44^

Here, we use a data-driven mechanistic modeling approach to elucidate the mechanisms of global human genome organization. In addition to accounting for the polymeric nature of individual chromosomes, we include particle-based representations for nucleoli, speckles, and nuclear lamina. Interactions within and among various components of the nucleus describe a coupled phase separation model for the diploid genome and nuclear bodies. After parameterization with Hi-C data, the nucleus model enables molecular dynamics simulations of genome structure and dynamics. Simulated 3D structures reproduce large-scale features of Hi-C data, correlate strongly with Lamin-B1 DamID and SON TSA-Seq, and match well with single-cell multiplexed genome imaging data. The co-phase separation model further captures the heterogeneous organization while producing well-defined distances between speckles and euchromatin. Speckles form through nucleation on chromatin segments, giving rise to close contacts that are preserved for a long time due to slow chromosome dynamics; given the non-specificity of phase separation, different sets of chromatin segments may nucleate speckle formation in different cells. Such heterogeneous contacts could further drive variations in chromosome positions. Together, our study highlights the significant impact of nuclear bodies on genome structure and dynamics.

## Results

### A co-phase separation model for the genome and nuclear bodies

Polymer simulations are useful tools for the mechanistic exploration of genome organization.^45–48^ They have been crucial for revealing the role of phase separation and loop extrusion in nuclear organization.^14,15,28,30,31,41,49–58^ In particular, the data-driven mechanistic modeling approach introduced in Ref. 33 directly links the quality of simulated genome structures with an energy function designed based on specific mechanisms of genome organization. As Hi-C data can constrain parameters of the energy function, the model’s performance in reproducing experimental data will be mainly determined by the quality of mechanistic assumptions rather than the uncertainty of parameters. Such a strategy is valuable for screening hypotheses and identifying mechanisms of global genome organization.

We generalize the data-driven mechanistic modeling approach to simulate the human nucleus. Each of the 46 chromosomes was represented as a beads-on-a-string polymer, with each bead corresponding to a one-MB-long genomic segment (see Figure 1). We labeled each chromatin bead as compartment *A, B*, or *C* for euchromatin, heterochromatin, or pericentromeric regions. Compartment-type specific interactions were incorporated to promote their phase separation. In addition to the block co-polymer setup, we introduced intrachromosome interactions that vary as a function of the sequence separation between two genomic segments. This “ideal” potential accounts for the role of specific protein molecules, such as loop extrusion via Cohesin molecules,^14,15^ and drives chromosome territory formation.^29,59^ Interactions among chromatin beads were optimized to reproduce various average contact probabilities determined from Hi-C experiments for HFF cells using the maximumentropy optimization algorithm. ^60–62^

**Figure 1:**
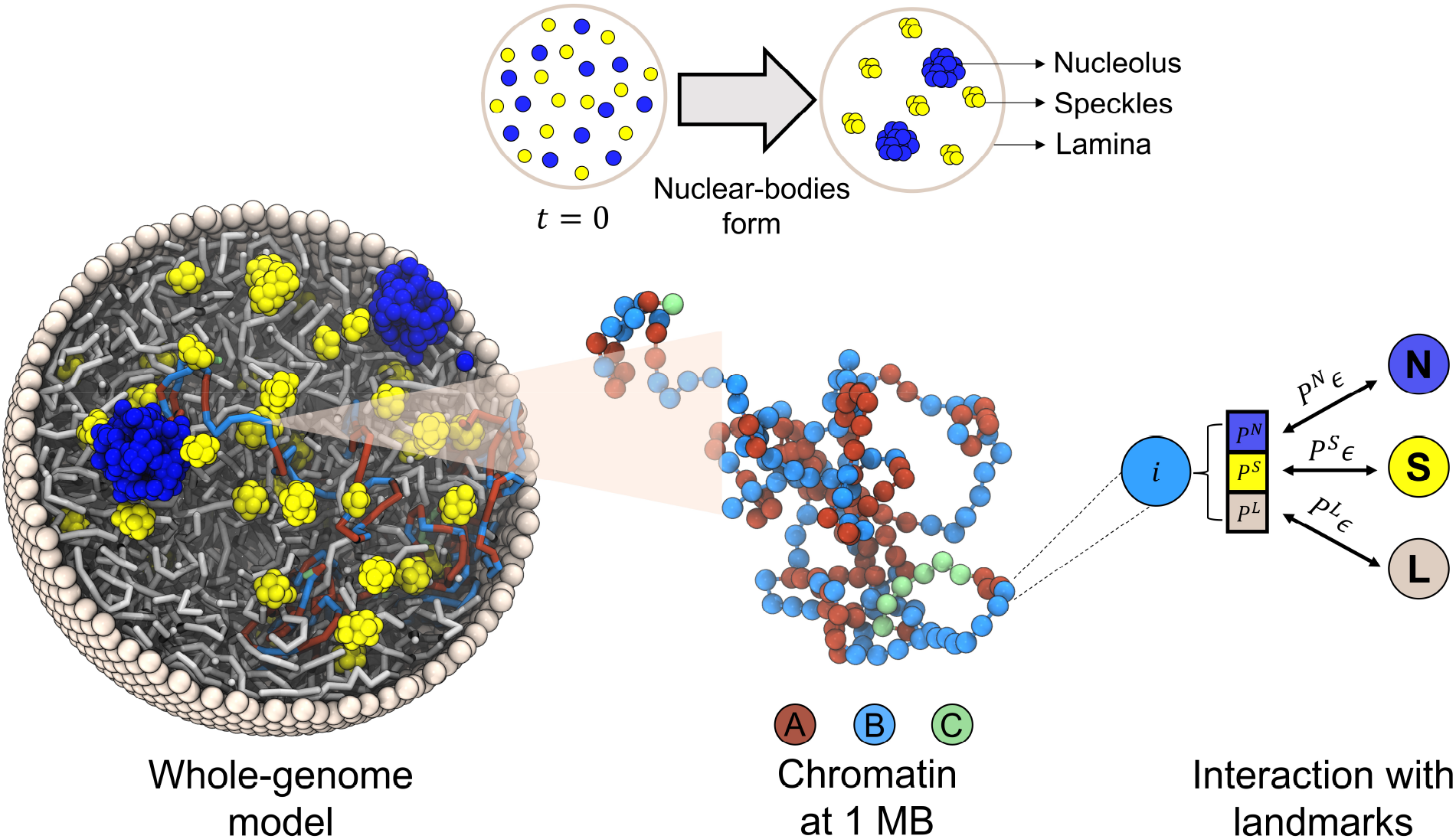
Illustration of the whole-genome model that explicitly considers chromosomes and various nuclear landmarks, including lamina (grey), nucleoli (blue) and speckles (yellow). Chromosomes are modeled as beads-on-a-string polymers at 1 MB resolution where the beads are further classified into euchromatin (red, compartment *A*), heterochromatin (light blue, compartment *B*) and centromeric regions (green, compartment *C*). As shown by the schematic in the upper panel, nucleoli and speckles form through the self assembly of coarse-grained particles uniformly distributed inside the nucleus at the beginning of simulations. Coupling between chromosomes and nuclear landmarks is accounted for with specific interactions, the strength of which depends on the contact probabilities (*P*^*N*^, *P*^*S*^, and *P*^*L*^) between them quantified using high-throughput sequencing data (see text).

In addition to the diploid genome, we adopted particle-based representation for various nuclear landmarks. We approximated the lamina as a spherical enclosure with a 10*µ*m diameter using discrete particles placed on a Fibonacci grid. The dynamics of the nuclear envelope was not considered, and the particles were fixed during simulations. Both nucleoli and speckles were modeled as liquid droplets^38,63^ that form through spontaneous phase separation of coarse-grained particles. These particles represent protein and RNA molecule aggregates and share attractive interactions within the same type to promote condensation. Coupling between chromatin and nuclear landmarks was accounted for with weak attractive interactions. Since their contacts are mediated by specific protein molecules, not all chromatin will form favorable contacts with every nuclear landmark. To ensure specificity, we further rescaled the strength of these interactions depending on the *intrinsic state* of chromatin beads. Three states, including speckle, nucleolus, and lamina state corresponding to the respective nuclear landmarks, were defined using SPIN.^64^ SPIN annotates chromatin based on their relative position with respect to various nuclear structures through an integrative analysis of TSA-Seq, DamID with Hi-C data. ^64^ We combined the 25-kb-resolution SPIN annotations to assign each chromatin bead with three probabilities for being in each state. These probabilities estimate the fraction of various states in the one-MB-long regions and were used to renormalize chromatin-nuclear landmark interactions.

We tuned the interaction parameters for nuclear particles to reproduce the corresponding nuclear bodies’ number and maximize the agreement between simulated and experimental DamID and TSA-Seq data. Attractive interactions between nucleolar particles and chromatin (see below) are sufficient to produce 2∼3 nucleoli.^65^ The chromatin network can nucleate phase separation and arrest the system in multi-droplet states. However, similar treatment for speckles failed to produce experimental numbers on the order of 30 ∼ 40 (see Figure S1). Instead, we introduced repulsive interactions in the form of the Yukawa potential in addition to attractive interactions among speckle particles. The Yukawa potential has been widely used for modeling colloids and is known to stabilize the multiple droplet state. ^66^ It can serve as an effective approximation to account for the impact of non-equilibrium modification to protein molecules that disrupts droplet coarsening.^67,68^ Detailed expression of the energy function and parameter values can be found in the Supporting Information.

### Validating model against sequencing data

Our computational model was designed with the assumption that the genome and nuclear bodies form through self-assembly during the early G1 phase. In particular, the euchromatin and heterochromatin phase separate with the presence of nucleolar and speckle particles that themselves organize into liquid droplets. However, the validity of our assumption depends on whether the co-phase separation model between chromosomes and nuclear bodies can produce structures that capture various features of nuclear organization. Upon parameterizing the interaction potentials, we extensively validated the simulated chromosome structures and chromosome-nuclear landmark contacts against experimental data to evaluate our model.

We carried out molecular dynamics simulations of the self-assembly process that organizes genome structures and drives the formation of nuclear bodies. A total of 100 independent, 12 million-timestep-long trajectories were simulated to yield an ensemble of 3D structures. The trajectories were initialized with chromosome configurations obtained from a separate sampling of a genome model introduced in a previous study. ^33^ Nucleolar and speckle particles were randomly distributed throughout the nucleus. The initial period of the simulations (6 million time steps) corresponds to a maturation process during which nucleoli and speckles form through spontaneous phase separation. While the dynamics of the maturation process is of interest, we did not include them for structural analysis.

We first examined the contacts between simulated chromosome structures and compared those with Hi-C experiments. As shown in Figure 2, the chromosomes form territories and occupy spatially distinct regions. ^22^ *A/B* compartments phase separate from each other, with *B* compartments preferentially localizing towards the nuclear periphery. The compartmentalization is evident both at the genome-wide scale and for individual chromosomes. Genome compartmentalization can also be seen from the checkerboard pattern of the simulated contact map (Figure 3a), which matches well with Hi-C data (see also Figure S2). Additionally, we computed the average contact probability between pairs of chromosomes. The reasonable agreement between simulated and experimental values suggests that phase separation and contacts with nuclear bodies contribute to inter-chromosome interactions (Figure S2c). The remaining discrepancy potentially arises from chromosome-specific interactions that the model does not capture.

**Figure 2:**
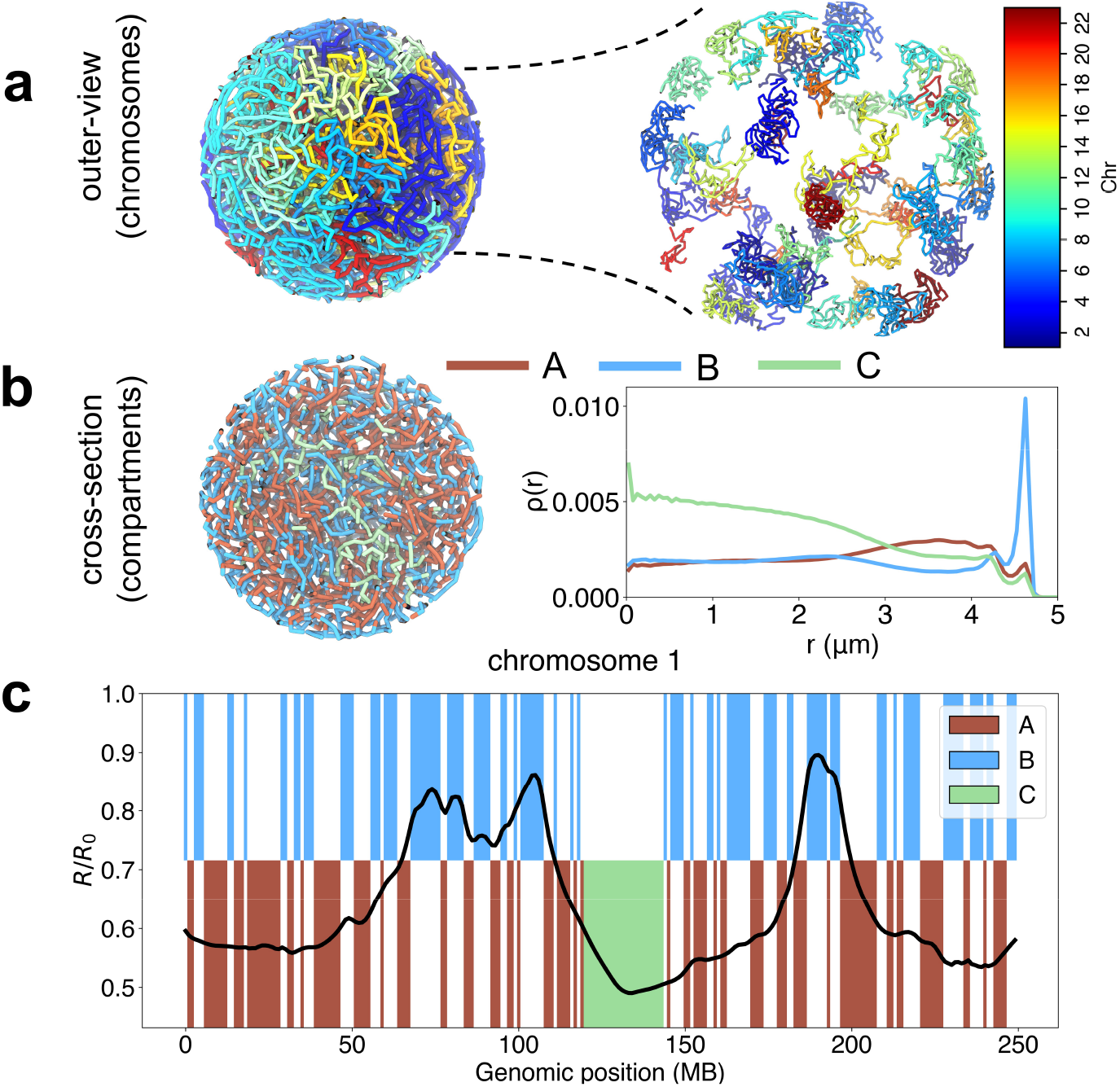
Simulated genome structures support chromosome territory formation and phase separation between euchromatin and heterochromatin. (a) Outer-view of the genome colored by chromosomes (homologs have the same color) and a blown-up version. (b) A cross-section of the nucleus is shown with chromatin colored by the compartment type, with *A* in red, *B* in blue, and *C* in green. The radial density of profiles of the three compartment types are shown on the side. (c) The average radial position of chromosome 1 as a function of the genomic position with comparison to the compartment profile.

**Figure 3:**
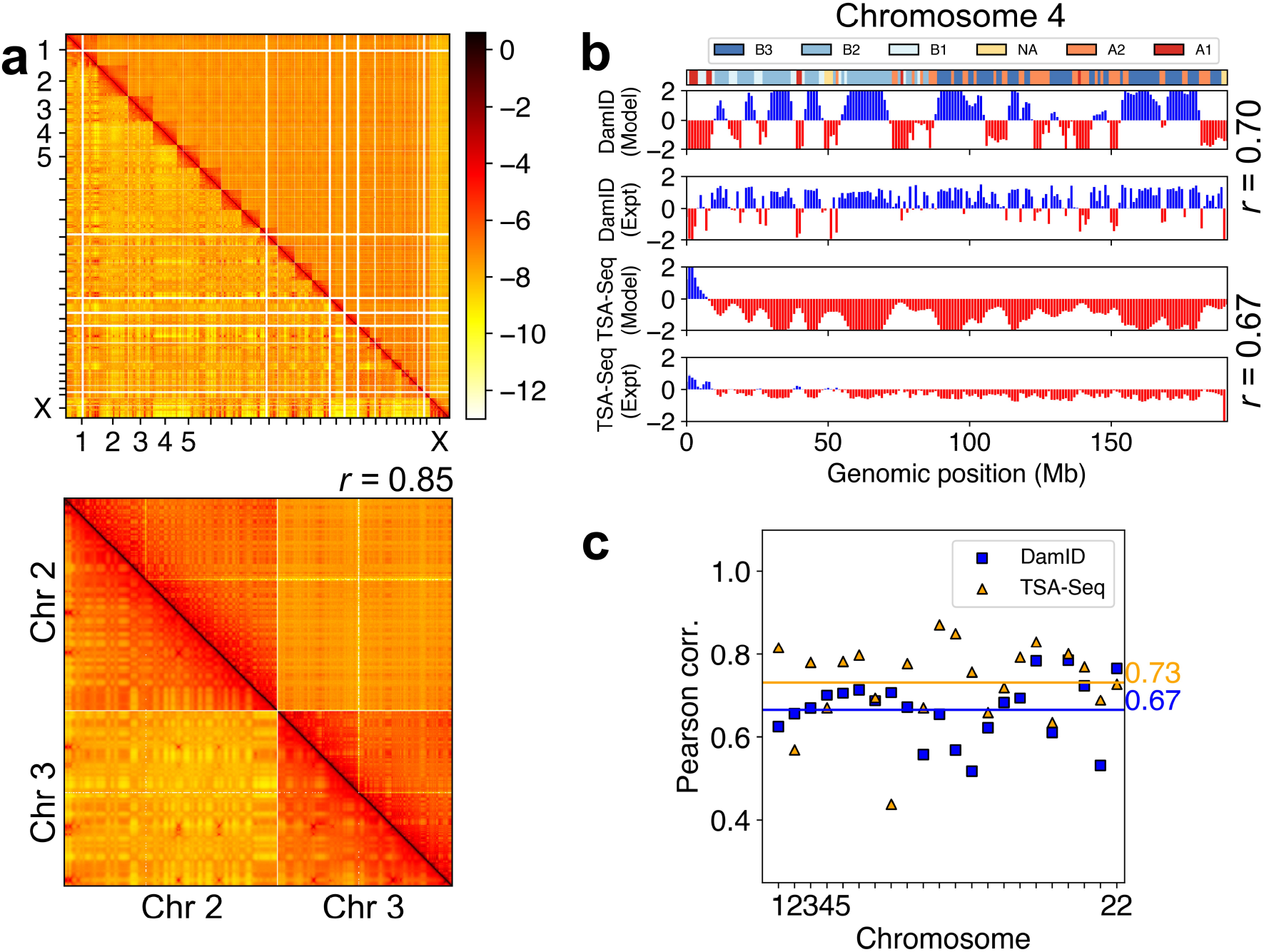
Simulated structures reproduce contacts within DNA segments and interactions between chromosomes and nuclear bodies. (a) Comparison between simulated (lower-triangle) and experimental (upper-triangle) contact matrix. ^69^ A zoomed-in contact map between chromosomes 2 and 3 is shown with the Pearson’s correlation coefficient (*r*). (b) Comparison (for chromosome 4) between the model predicted Lamin-B DamID and SON TSA-Seq signals and the experimental data at 1MB resolution, with the Pearson’s correlation coefficients shown on the side. The subcompartment annotations at the top of the figure were taken from the IMR90 cell type (fibroblast). ^70^ (c) Pearson correlation coefficients between simulated and experimental Lamin-B DamID (blue) and SON TSA-Seq (yellow) data for individual chromosomes. The genome-wide averages are shown as straight lines with the corresponding values on the side.

We computed chromatin-speckle and chromatin-lamina contacts to yield *in silico* predictions of the SON TSA-Seq and Lamin-B DamID signals, respectively. The simulated and the experimental SON TSA-Seq signals^23^ are in good agreement with a consistent trend, as shown in the bottom two panels of Figure 3b for chromosome 4. The Pearson correlation coefficient between the two is 0.67 (chr 4), and the genome-wide average correlation score is 0.73 (Figure 3c). The DamID signals are also reproduced satisfactorily, with a genome-wide correlation score of 0.67. We note that the 1Mb resolution used in our model may be insufficient to capture the sharp transitions between contact-enriched (shown in blue in Fig 3b) and contact-depleted zones (shown in red) with short periods, preventing a perfect reproduction of experimental data. Nevertheless, as shown in the following sections, the model’s computational efficiency allows comprehensive mechanistic exploration of the coupling between chromatin organization and nuclear landmarks.

### Validating model against imaging data

In addition to sequencing-based genomic mapping data, we utilized the multiplexed 3D genome imaging data to further evaluate our model. Recent advancements in microscopy imaging-based techniques have enabled simultaneous detection of hundreds to thousands distinct genomic loci.^25,71^ These imaging data provide valuable information on the 3D positions of genomic regions at single-cell resolution for more direct comparisons with the simulated structures in order to benchmark the quality of our model.

We first examined that in our simulations, the nuclear bodies formed through self-assembly of individual particles that are randomly distributed at the start. As shown in Figure S3, the average number of nucleoli (2) and speckles (36) are in good agreement with values estimated from microscopic images.^73,74^ While the classical nucleation theory would predict an equilibrium state with single droplets, the attractive interactions between nucle-oli and chromatin help arrest droplet coalescence, stabilizing the multi-droplet state. ^65^ The weak but long-ranged repulsive interactions in addition to the short-ranged attractive interaction among the speckle particles further suppress droplet coarsening, producing an order of magnitude more speckles than nucleoli.

We further evaluated chromosome conformations against imaging data. As shown in Figure 4, the simulated radial chromosome positions are highly correlated with experimental values^24^ with a Pearson correlation coefficient of 0.68. The radius of gyration of individual chromosomes matches well with DNA-MERFISH imaging data that report the spatial positions of uniformly selected loci across chromosomes in individual cells.^25^

**Figure 4:**
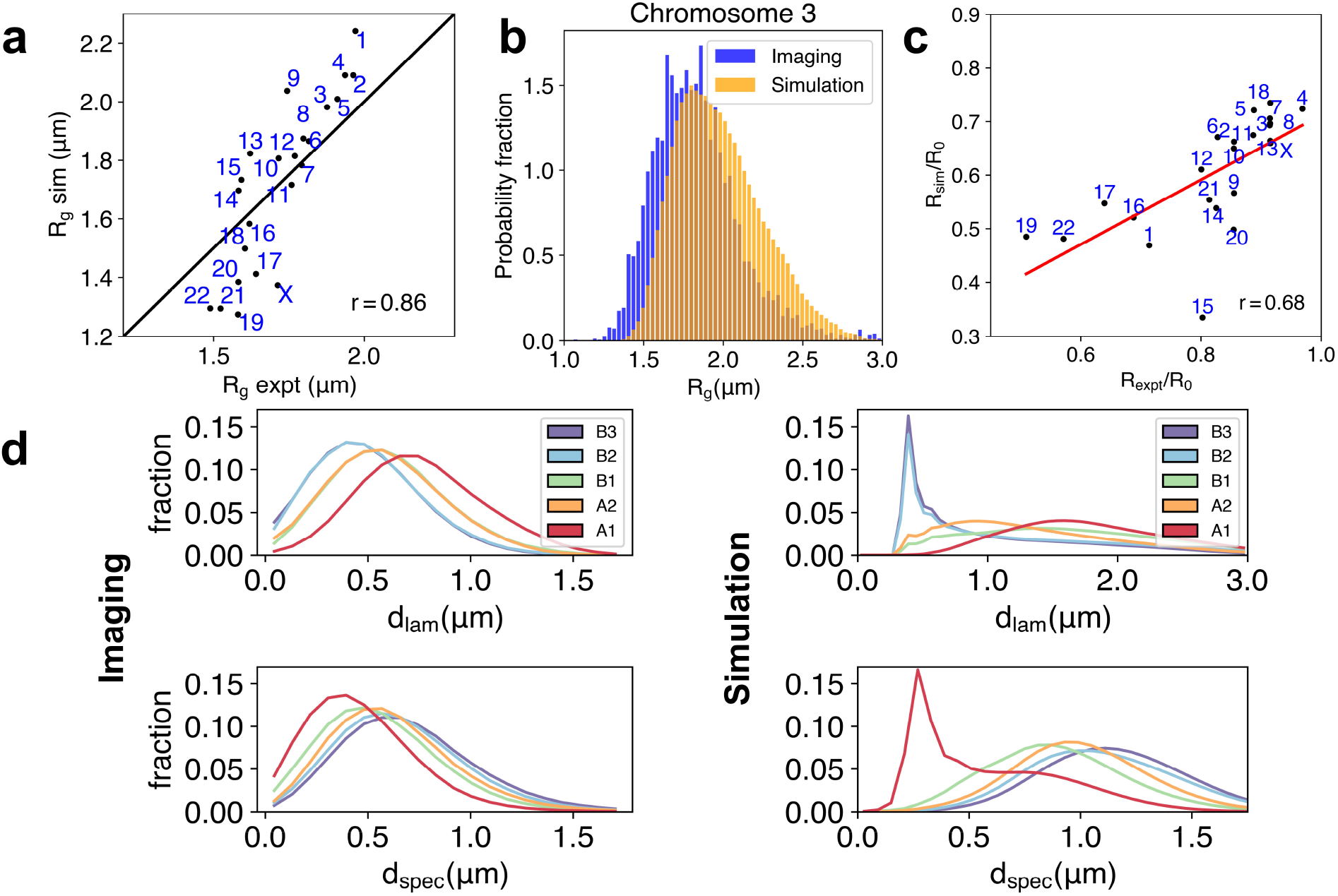
Simulated structures reproduce chromosome sizes, positions, and the localization of genome subcompartments. (a) Comparison between chromosome radius of gyrations from simulations and from DNA-MERFISH experiments. ^25^ The Pearson’s correlation coefficient (*r* = 0.86) is also shown. (b) Overlap of the probability distributions for the radius of gyration of chromosome 3 computed from the experimental and simulated structural ensemble. (c) Comparison of the chromosome radial positions in experiment^72^ and simulations. (d) Distribution of genome subcompartments as a function of distance from nuclear lamina (*d*_lam_) and speckles (*d*_spec_), respectively, shown for DNA-MERFISH imaging^25^ and simulation data.

Finally, we characterized the localization of specific genomic regions relative to nuclear bodies. In particular, we study the five primary subcompartments that further divide euchromatin and heterochromatin into *A*1, *A*2, *B*1, *B*2, and *B*3.^11^ Subcompartments provide a more nuanced classification of chromatin than compartments *A/B* to distinguish the varying degrees of gene activation/repression and contacts with nuclear landmarks.^36^ We found that the simulated contacts are in qualitative agreement with DNA-MERFISH imaging results (Figure 4). For example, subcompartment *A*1 and *B*3 strongly localize in close proximity to speckles and nuclear lamina respectively. Additionlly, subcompartments *A*2 and *B*2 occupy intermediate regions transitioning between speckles and nuclear lamina. *A*2 is localized more towards speckles, while *B*2 is close to the nuclear lamina. We note that a quantitative comparison with imaging data was not attempted because we used a spherical model for the nuclear lamina, while cells in experiments are more ellipsoidal. ^25^

It is worth pointing out that our model only recognizes *A* and *B* compartments but was not explicitly provided with the subcompartment annotations. Its success in differentiating the spatial localization of subcompartments arises from the chromatin-nuclear landmark interactions encoded with SPIN states. For example, *A*1 and *B*3 strongly correlate with speckle and lamina states, ^64^ resulting in their preferential contacts with the respective nuclear landmarks seen in Figure 4. Together, SPIN states and compartment types provide complementary representations of the genome to capture the distinct aspects of nuclear organization.

### Interactions with speckles refine *A* compartments

Our extensive validations support the co-phase separation model for chromosome organization and nuclear body formation. The coupling between chromatin and nuclear lamina helps position heterochromatin towards the nuclear periphery. Without such coupling, we previously showed that heterochromatin would occupy interior locations.^33^ The impact of speckles on euchromatin is less clear since the phase separation between euchromatin and heterochromatin will naturally place them towards nuclear interior. Whether speckles reinforce euchromatin’s interior position remains to be seen.

To probe the impact of speckles on genome organization, we perturbed the system to induce speckle coalescence by weakening the repulsive potential between speckle particles (Figure 5). Fusing droplets could bring associated chromatin segments together, exerting a pulling force on the genome, as shown by Brangwynne and coworkers. ^75^ We found that the spatial distribution of *A* compartments was altered significantly as speckle numbers decreased from 40 to 10. Speckles and euchromatin tend to shift towards the nuclear interior to maximize contacts with *A* compartments coming from all directions upon coalescence. A similar movement of speckles has indeed been seen in cells with inhibited transcription that induce speckle fusion.^76^ We further broke down the impact into two subcompartments. As expected, there is a significant perturbation to *A*1 that directly interact with speckles. Notably, we found the distribution of *A*2 is similarly altered as well, though to a less extent. Since most of *A*2 do not directly contact speckles, their impact is an indirect effect mediated via *A*1 as a result of the attraction among all *A* compartments.

**Figure 5:**
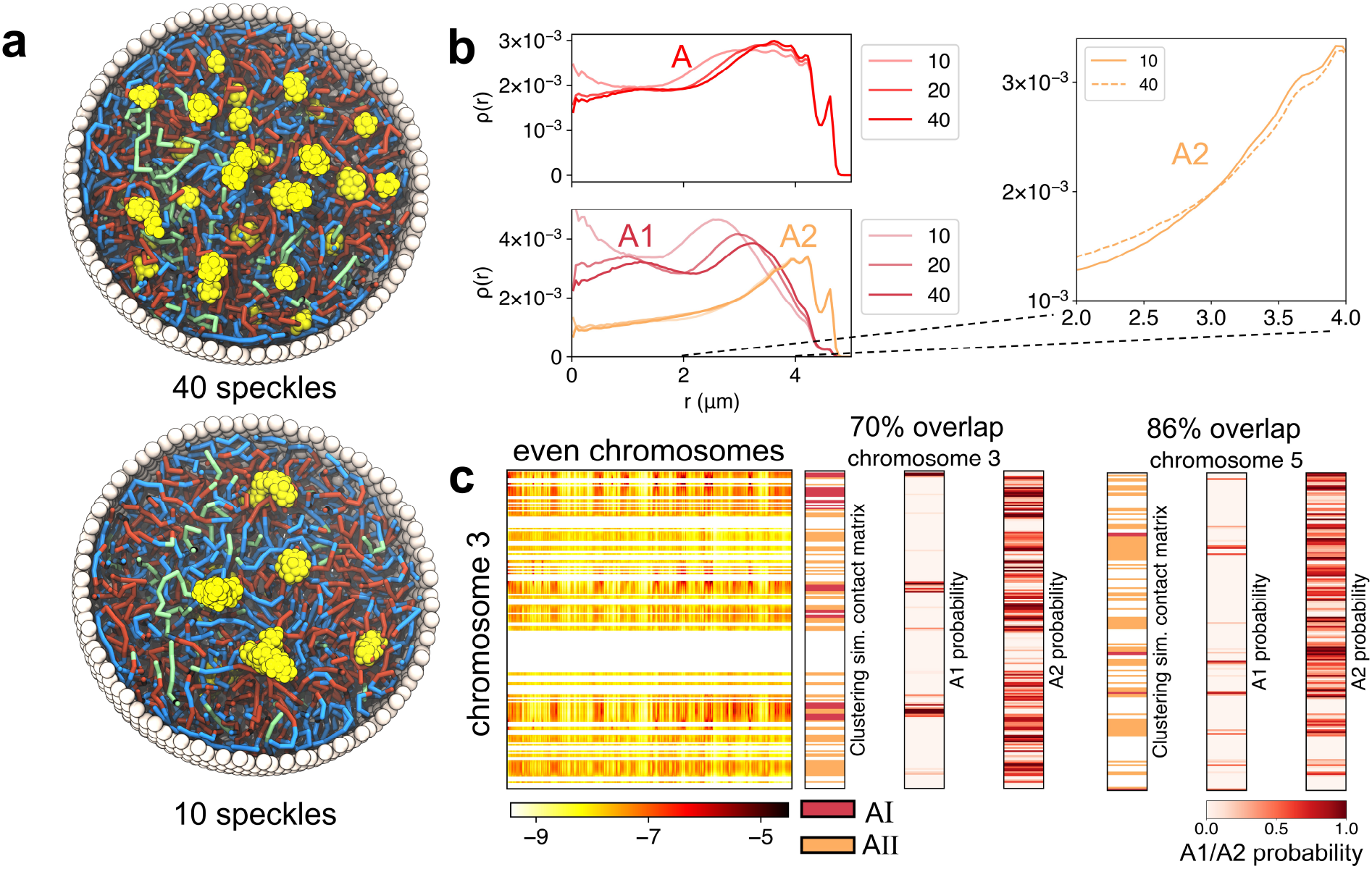
Speckles pull a subset of *A* compartments towards nuclear interior, producing their distinct contact patterns with the rest of the genome. (a) Cross sections of two nucleus models with 40 (top) and 10 (bottom) speckles. Speckles in the bottom panel adopt more interior localizations. (b) Radial densities of the *A* compartment and *A*1, *A*2 subcompartments as a function of the radial position inside the nucleus. The upper panel highlights the pulling of the *A* segments to the interior on lowering the number of speckles in the system. The lower panel highlights the behavior of the *A*1 and *A*2 on perturbing the number of speckles in the system. (c) Comparison between the two sub-types of *A* compartments (*AI, AII*) and *A*1 and *A*2 subcompartment annotation probabilities for chromosomes 3 and 5. The simulated genome-wide contact matrix between *A* loci on chromosome 3 and all even chromosomes is shown on the left. This contact matrix was used for clustering the *A* loci into two sub-types (see also Figure S4).

The correlation between the speckle state and the subcompartment *A*1 drives the coupling between speckle fusion and chromatin interior movement. Given the correlation, it is tempting to hypothesize that the partition of compartment *A* into two subcompartments results from their differential contacts with nuclear speckles. This hypothesis implies that interacting with speckles alters the contact patterns between the subset of *A* compartments with the rest of the genome. As a test of this hypothesis, we performed a Gaussian hidden Markov model clustering of all *A* compartment beads based on their inter-chromosomal contact patterns. The same procedure introduced in Rao et al.^11^ that led to the annotation of subcompartments was adopted here. As shown in Figure 5 and Figure S4, the two clusters identified from our simulated contact map indeed correlate well with *A*1*/A*2 annotations based on the Hi-C data. Therefore, differential contacts with speckles may underly the unique contact patterns seen in Hi-C data and the rise of subcompartments. Our results support that nuclear bodies and self-organization play an essential role in refining the compartments.

### Single cell heterogeneity and robustness

The results presented so far support that the model captures the average contacts within chromatin and between chromatin and nuclear landmarks. Next, we analyze the simulated structural ensemble to characterize the heterogeneity and fluctuation of these contacts. Specifically, we sought to ask whether the model can reconcile the apparent heterogeneity of genome organization across cells with the emergence of well-defined distances, which may appear paradoxical as discussed in the *Introduction*.

To quantify the heterogeneity of genome organization, we computed the variance of chromosome positions and chromosome radius of gyration across a total of 100 trajectories that are initialized from distinct configurations (see Methods). As shown in Figure S5, chromosome position distributions from independent simulations differ significantly, with a distribution width of about 1.4 *µ*m. Chromosome radii of gyration also undergo substantial changes (Figure 4 and Figure S6), on an order that quantitatively agrees with values from DNA-MERFISH imaging.^25^

In addition to the heterogeneity at chromosome levels, we further studied local chromatin organization around speckles. We computed the average contact map for *A*1 subcompartments that are known to localize close to speckles.^36^ Two *A*1 segments were noted as in contact if they are bound to the same speckle. Figure 6 shows the variance of the speckle mediated *A*1 contact map across simulation trajectories. There is a significant fluctuation, consistent with the variation of chromosome positions shown in Figure S5. This variation arises from the non-specificity of phase separation: while *A*1 segments almost always interact with the speckle particles and nucleate their condensation, different sets of *A*1 may contribute to such interactions in different trajectories.

**Figure 6:**
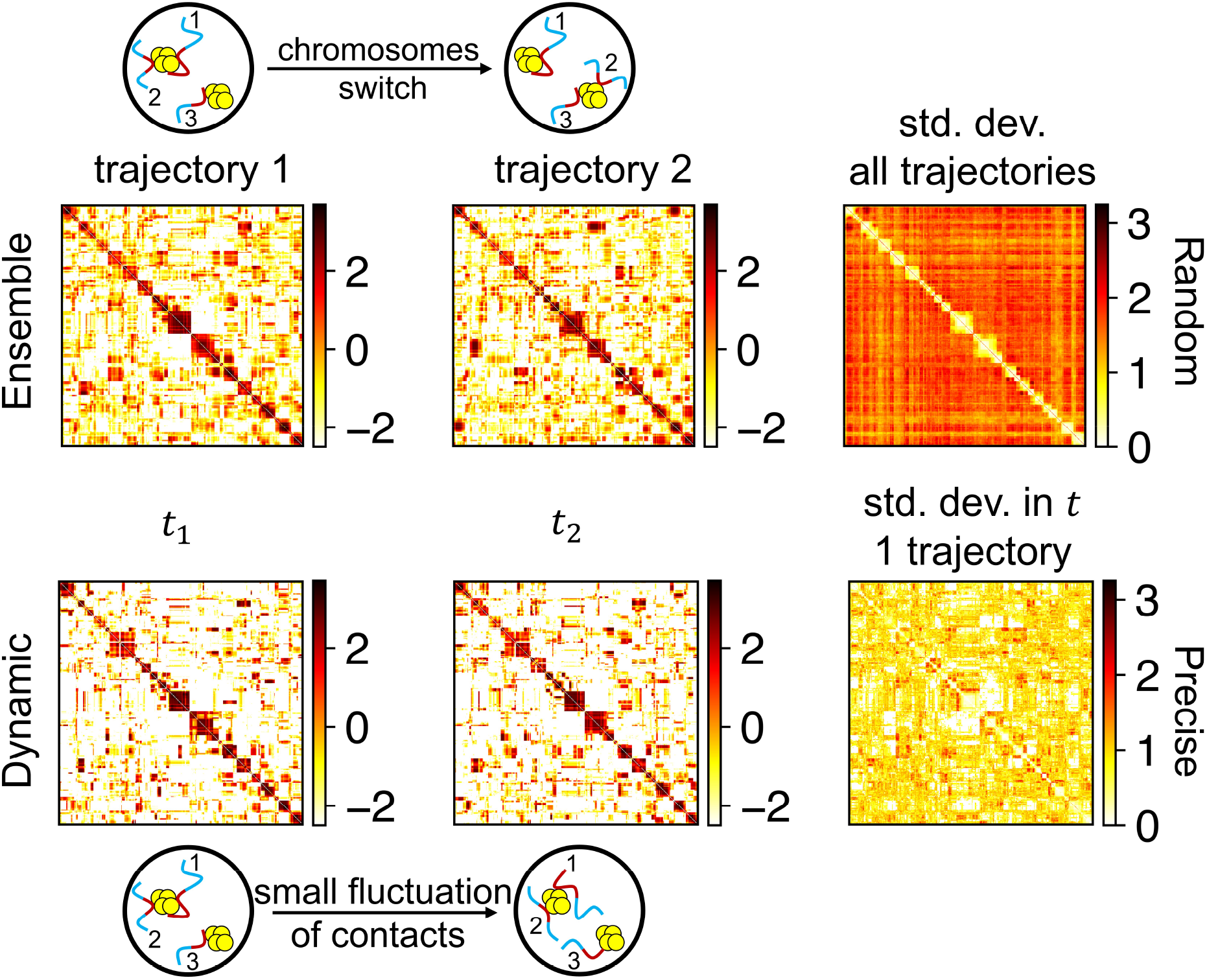
Variation of speckle-mediated contacts between genomic segments of *A*1 subcompartments. The upper panel shows the average *A*1 − *A*1 contact matrix computed from independent simulation trajectories, and the lower panel shows the contact matrix calculated at different times within the same trajectory. The matrix on the far right in both panels denotes the standard deviation in the contact matrices. The dissimilarity in *A*1 contacts between different trajectories has a much higher variation as compared to the variance in the contacts within a trajectory.

In contrast to the large fluctuations across independent trajectories, we found that the contacts among *A*1 evaluated at different time points of the same trajectory are relatively robust with minimal changes. Minimal variations within one trajectory arise since speckle chromatin interactions are strong and can last for a long time. The *A*1 segments are essentially glued with their *A*1 counterparts formed at the beginning of phase separation. Therefore, the co-phase separation model, in which speckles nucleate around chromatin segments, captures both the random and precise nature of genome organization.

## Discussion

In this work, we developed a molecular simulation framework to study the human genome organization and explore mechanisms for its setup. In addition to polymer models of chromosomes, we explicitly incorporated particle-based representations of nuclear lamina, nucleoli, and speckles. The 3D organization of chromosomes and the formation of nuclear bodies were modeled as a self-assembly process driven by the various interaction energies in the system. Such self-assembly simulations reproduced global features of genome structures, the number of nuclear bodies, and the contacts between the genome and nuclear landmarks. Our results support the hypothesis that nuclear landmarks largely compartmentalize the genome by attracting active and repressive regions to the nuclear interior and periphery respectively. The formation of speckles that strongly interact with chromatin introduces effective crosslinks to the polymeric chromatin network. A cross-linked system is consistent with the slow dynamics of speckles^76,77^ and gel-like behavior of chromatin.^78,79^ Such a cross-linked system formed through phase separation provides an intuitive explanation for the random yet precise global genome organization discussed in the *Introduction* section. Nuclear bodies form through nucleation on highly transcribed chromatin segments. This nucleation mainly occurs during the early G1 phase and may encode an initial memory of speckle-chromatin contacts. The cross-linking and tethering by nuclear lamina constrains chromosome dynamics to preserve the memory, producing well-defined distances between genome and nuclear bodies. Slow chromatin dynamics is indeed well documented in the literature,^78–81^ potentially as a result of interactions with nuclear bodies.^82^ On the other hand, because of the non-specificity of phase separation, different sets of chromatin segments could participate in different cells to nucleate speckle formation. This heterogeneity of contacts across cells could further drive the fluctuation of chromosome radial positions. While heterogeneous, the co-phase separation model does ensure that genomic regions that are highly transcribed will always reside in spatial proximity with speckles. Notably, encoding the well-defined, functionally important distances does not require the system to reach equilibrium, which can be challenging given the genome’s large size. Further exploring genome dynamics with the model introduced here will represent exciting future directions.

We note that in addition to the nuclear landmarks considered here, other molecules, including Lamin A^83^ and HP1,^84^ have also been suggested to crosslink chromatin. The coarsened resolution of our model would not be sufficient to capture the impact of these molecules, which may work together with speckles to constrain the genome structure and dynamics further.

Several studies have recently attempted to reconstruct whole-genome organizations from experimental data. ^32,85^ Our study is complementary to the integrative modeling approach by Boninsegna et al. ^85^, which combined Hi-C, Lamin-B1 DamID, 3D FISH imaging, and SPRITE data to produce an ensemble of genome structures. The modeled structures were shown to predict orthogonal experimental data from SON TSA-seq and DNA-MERFISH imaging well. Like the study of Boninsegna et al. ^85^, our approach succeeded in building consensus structures that agree with a collection of experimental data. However, the explicit modeling of nuclear landmarks and their dynamical coupling with chromatin that are unique to this study render our approach useful for uncovering mechanisms that lead to the establishment of such structures. The model introduced by Fujishiro and Sasai ^32^ is more similar in spirit to our approach by designing an effective energy function of polymer models that can reproduce/explain experimental data. However, they did not consider the role of speckles in genome organization. As discussed in the main text, speckles are essential for an accurate description of the structure and dynamics of active chromatin.

## Methods

### Detailed setup of nucleus model

We modeled all 46 chromosomes of the diploid human genome at the 1 MB resolution using a total of 6106 coarse-grained beads. Each chromatin bead was assigned as compartment *A, B, C*, or *N*. *A/B* compartments were determined from the first eigenvectors of the intrachromosomal Hi-C contact matrices, and compartment *C* were identified as centromeric regions based on the DNA sequence. Compartment *N* denote genomic regions that cannot be assigned as either *A, B*, or *C* due to a lack of Hi-C data. No interaction parameters were assigned for compartment *N*. In addition to the compartment assignments, each chromatin bead was provided with three probabilities (*P*^N^, *P*^S^, and *P*^L^) that denote their tendency to interact with the three nuclear landmarks. More details on computing these probabilities are provided in the Supplementary Information. Based on super-resolution imaging data, ^86^ we estimated the size of each bead as 385 nm.

The number of coarse-grained particles for nucleoli (300) and speckles (600) was estimated based on the experimentally reported values of nuclear protein NPM1 concentration as done in a recent study^65^ and the protein densities calculated from refractive index measurements.^87^ The size of these particles was estimated as 192.5 nm based on the average radius of individual nucleoli and speckles. We note that the above estimation is crude. The nucleolar and speckle particles should be viewed as molecular aggregates rather than a single protein molecule. Given the size of a typical protein as 5 ∼ 10 *nm*,^88^ the number of molecules within a single coarse-grained particle can be on the order of 10^3^. This number, while large, is in the same order as the number of distinct molecules that make up the nucleoli.^89^

### Molecular dynamics simulation details

We used the software package LAMMPS^90^ to perform molecular dynamics (MD) simulations in reduced units. Constant-temperature (*T* = 1.0 in reduced unit) simulations were carried out via the Langevin dynamics with a damping coefficient *γ* = 10.0 and a simulation time step of *dt* = 0.005. We froze the lamina particles and only propagated the dynamics of chromatin, nucleoli and speckles. Configurations were recorded at every 2000 simulation steps for analysis. The initial configurations of independent MD simulations were built as follows. We first obtained whole-genome structures by uniform sampling from a 20-million-step long trajectory carried out in our previous study. ^33^ This trajectory was obtained without constraints of nuclear bodies and captures the large fluctuation of chromosome positions. Next, 300 nucleoli and 600 speckle particles were placed with random positions inside the nucleus. We ran a short minimization to remove any potential overlaps before carrying out each simulation for a total of twelve million timesteps. The first six million time steps of each trajectory were discarded as equilibration.

### Data processing and analysis

#### Experimental data

We obtained the *in situ* Hi-C data of HFF cell lines from the 4DN data portal. The intra- and inter-chromosomal interactions were calculated at 1Mb resolution with VC SQRT normalization applied to the interaction matrices. Hi-C data extraction and normalization were performed using Juicer tools. ^91^ For Hi-C subcompartments, we used the IMR90 Hi-C sub-compartment annotation produced by SNIPER.^70^ SON TSA-seq data in HFF cell line is obtained from the 4DN data portal. The TSA-seq processing and normalization method is described in Ref. 23. Lamin-B DamID data in HFF cell line is obtained from the 4DN data portal. Two biological replicates were merged and the normalized counts over Dam-only control were used for analysis.

The SON TSA-Seq and Lamin-B DamID data were processed at the 25 KB resolution and the average values at the 1 MB resolution were used in Figure 3 for model validation. As the subcompartment annotation was performed at the 100 kb resolution, we assigned each 1MB bead in our model with probabilities for being in various subcompartments estimated from arithmetic means. We further assigned beads as *A*1 when computing Figure 6 if the corresponding probabilities are higher than 0.5.

#### Simulation data

Complete details on the analysis of simulated structures can be found in the SI, and here we briefly describe the procedure. To compare with SON TSA-Seq data, we used the DBSCAN algorithm^92^ to identify speckle clusters in our simulations, and consecutively, the distances of the speckles from different chromatin segments. We calculated the *in silico* SON TSA-Seq signal for a chromatin segment using a speckle distance-dependent exponential decay function commensurate with the TSA-Seq microscopy experiments. ^36^ The signals were then converted into enrichment scores after calculating the genome-wide averages of TSA-Seq signals.

For the comparison with Lamin-B DamID, we used the radial positions of chromatin segments to calculate the distance from the lamina. If the chromatin segment is within a threshold distance of the lamina, we identified the two as in contact. We calculated the *in silico* DamID signals as the contact probabilities across all simulation trajectories.

## Acknowledgement

This work was supported in part by the National Institutes of Health grant R35GM133580 (B.Z.) and National Institutes of Health Common Fund 4D Nucleome Program grant UM1HG011593 (J.M.). We thank Bas van Steensel for making the LaminB DamID data for HFF cells available.

## Author Contributions

Conceptualization: KK, BZ

Methodology: KK, YFQ, BZ

Investigation: KK, YFQ, YCW, JM, BZ

Visualization: KK, BZ Supervision: BZ

Writing original draft: KK, BZ

Writing review & editing: KK, YFQ, YCW, JM, BZ

## Data Availability

Hi-C data (https://data.4dnucleome.org, accession number: 4DNES2R6PUEK). Hi-C subcompartments (https://cmu.app.box.com/s/n4jh3utmitzl88264s8bzsfcjhqnhaa0/folder/86847603885). SON TSA-seq data (https://data.4dnucleome.org, accession number: pulldown data 4DNEX6U8TS3Y, control data 4DNEXI7XUWFK). LaminB DamID data (https://data.4dnucleome.org, accession number 4DNESXZ4FW4T).

## References

(1) Hübner, M. R.; Eckersley-Maslin, M. A.; Spector, D. L. Chromatin organization and transcriptional regulation. Curr. Opin. Genet. Dev. 2013, 23, 89–95.

(2) Bickmore, W. A. The Spatial Organization of the Human Genome. Annu. Rev. Genomics Hum. Genet. 2013, 14, 67–84.

(3) Gorkin, D. U.; Leung, D.; Ren, B. The 3D genome in transcriptional regulation and pluripotency. Cell Stem Cell 2014, 14, 762–775.

(4) Cremer, T.; Cremer, M.; Hubner, B.; Strickfaden, H.; Smeets, D.; Popken, J.; Sterr, M.; Markaki, Y.; Rippe, K.; Cremer, C. The 4D nucleome: Evidence for a dynamic nuclear landscape based on co-aligned active and inactive nuclear compartments. FEBS Lett 2015, 589, 2931–2943.

(5) Bonev, B.; Cavalli, G. Organization and function of the 3D genome. Nat. Rev. Genet. 2016, 17, 661–678.

(6) Dekker, J.; Mirny, L. The 3D Genome as Moderator of Chromosomal Communication. Cell 2016, 164, 1110–1121.

(7) Hnisz, D.; Day, D. S.; Young, R. A. Insulated Neighborhoods: Structural and Functional Units of Mammalian Gene Control. Cell 2016, 167, 1188–1200.

(8) Rowley, M. J.; Corces, V. G. Organizational principles of 3D genome architecture. Nat. Rev. Genet. 2018, 19, 789–800.

(9) Finn, E. H.; Misteli, T. Molecular basis and biological function of variability in spatial genome organization. Science 2019, 365, eaaw9498.

(10) Furlong, E. E.; Levine, M. Developmental enhancers and chromosome topology. Science 2018, 361, 1341–1345.

(11) Rao, S. S.; Huntley, M. H.; Durand, N. C.; Stamenova, E. K.; Bochkov, I. D.; Robinson, J. T.; Sanborn, A. L.; Machol, I.; Omer, A. D.; Lander, E. S.; Aiden, E. L. A 3D map of the human genome at kilobase resolution reveals principles of chromatin looping. Cell 2014, 159, 1665–1680.

(12) Nora, E. P.; Lajoie, B. R.; Schulz, E. G.; Giorgetti, L.; Okamoto, I.; Servant, N.; Piolot, T.; Van Berkum, N. L.; Meisig, J.; Sedat, J.; Gribnau, J.; Barillot, E.; Blüthgen, N.; Dekker, J.; Heard, E. Spatial partitioning of the regulatory landscape of the X-inactivation centre. Nature 2012, 485, 381–385.

(13) Dixon, J. R.; Selvaraj, S.; Yue, F.; Kim, A.; Li, Y.; Shen, Y.; Hu, M.; Liu, J. S.; Ren, B. Topological domains in mammalian genomes identified by analysis of chromatin interactions. Nature 2012, 485, 376–380.

(14) Sanborn, A. L. et al. Chromatin extrusion explains key features of loop and domain formation in wild-type and engineered genomes. Proceedings of the National Academy of Sciences 2015, 112, E6456–E6465.

(15) Fudenberg, G.; Imakaev, M.; Lu, C.; Goloborodko, A.; Abdennur, N.; Mirny, L. A. L. A. Formation of Chromosomal Domains by Loop Extrusion. Cell Rep 2016, 15, 2038–2049.

(16) Xie, W. J.; Qi, Y.; Zhang, B. Characterizing chromatin folding coordinate and landscape with deep learning. PLOS Comput. Biol. 2020, 16, e1008262.

(17) Parmar, J. J.; Woringer, M.; Zimmer, C. How the Genome Folds: The Biophysics of Four-Dimensional Chromatin Organization. Annu. Rev. Biophys. 2019, 48, 231–253.

(18) Fudenberg, G.; Abdennur, N.; Imakaev, M.; Goloborodko, A.; Mirny, L. A. Emerging Evidence of Chromosome Folding by Loop Extrusion. Cold Spring Harb. Symp. Quant. Biol. 2017, 82, 45–55.

(19) Williams, R. R.; Fisher, A. G. Chromosomes, positions please! Nat. Cell Biol. 2003, 5, 388–390.

(20) Parada, L. A.; Roix, J. J.; Misteli, T. An uncertainty principle in chromosome positioning. Trends Cell Biol. 2003, 13, 393–396.

(21) Bickmore, W. A.; Chubb, J. R. Chromosome position: Now, where was I? Curr. Biol. 2003, 13, R357–R359.

(22) Cremer, T.; Cremer, C. Chromosome territories, nuclear architecture and gene regulation in mammalian cells. Nat. Rev. Genet. 2001, 2, 292–301.

(23) Zhang, L.; Zhang, Y.; Chen, Y.; Gholamalamdari, O.; Wang, Y.; Ma, J.; Belmont, A. S. TSA-seq reveals a largely conserved genome organization relative to nuclear speckles with small position changes tightly correlated with gene expression changes. Genome Res. 2021, 31, 251–264.

(24) Boyle, S.; Gilchrist, S.; Bridger, J. M.; Mahy, N. L.; Ellis, J.; Bickmore, W. A. The spatial organization of human chromosomes within the nuclei of normal and emerinmutant cells. Hum. Mol. Genet. 2001, 10, 211–219.

(25) Su, J. H.; Zheng, P.; Kinrot, S. S.; Bintu, B.; Zhuang, X. Genome-Scale Imaging of the 3D Organization and Transcriptional Activity of Chromatin. Cell 2020, 182, 1641– 1659.e26.

(26) Payne, A. C.; Chiang, Z. D.; Reginato, P. L.; Mangiameli, S. M.; Murray, E. M.; Yao, C.-C.; Markoulaki, S.; Earl, A. S.; Labade, A. S.; Jaenisch, R., et al. In situ genome sequencing resolves DNA sequence and structure in intact biological samples. Science 2021, 371.

(27) Thomson, I.; Gilchrist, S.; Bickmore, W. A.; Chubb, J. R. The Radial Positioning of Chromatin Is Not Inherited through Mitosis but Is Established de Novo in Early G1. Curr. Biol. 2004, 14, 166–172.

(28) Jost, D.; Carrivain, P.; Cavalli, G.; Vaillant, C. Modeling epigenome folding: Formation and dynamics of topologically associated chromatin domains. Nucleic Acids Res. 2014, 42, 9553–9561.

(29) Di Pierro, M.; Zhang, B.; Aiden, E. L.; Wolynes, P. G.; Onuchic, J. N. Transferable model for chromosome architecture. Proc. Natl. Acad. Sci. 2016, 113, 12168–12173.

(30) Falk, M.; Feodorova, Y.; Naumova, N.; Imakaev, M.; Lajoie, B. R.; Leonhardt, H.; Joffe, B.; Dekker, J.; Fudenberg, G.; Solovei, I.; Mirny, L. A. Heterochromatin drives compartmentalization of inverted and conventional nuclei. Nature 2019, 570, 395–399.

(31) Shi, G.; Liu, L.; Hyeon, C.; Thirumalai, D. Interphase human chromosome exhibits out of equilibrium glassy dynamics. Nat. Commun. 2018, 9, 3161.

(32) Fujishiro, S.; Sasai, M. Generation of dynamic three-dimensional genome structure through phase separation of chromatin. bioRxiv 2021,

(33) Qi, Y.; Reyes, A.; Johnstone, S. E. S.; Aryee, M. M. J.; Bernstein, B. B. E.; Zhang, B. Data-Driven Polymer Model for Mechanistic Exploration of Diploid Genome Organization. Biophys. J. 2020, 119, 1905–1916.

(34) Pickersgill, H.; Kalverda, B.; De Wit, E.; Talhout, W.; Fornerod, M.; Van Steensel, B. Characterization of the Drosophila melanogaster genome at the nuclear lamina. Nat. Genet. 2006, 38, 1005–1014.

(35) Quinodoz, S. A. et al. Higher-Order Inter-chromosomal Hubs Shape 3D Genome Organization in the Nucleus. Cell 2018, 174, 744–757.e24.

(36) Chen, Y.; Zhang, Y.; Wang, Y.; Zhang, L.; Brinkman, E. K.; Adam, S. A.; Goldman, R.; Van Steensel, B.; Ma, J.; Belmont, A. S. Mapping 3D genome organization relative to nuclear compartments using TSA-Seq as a cytological ruler. J. Cell Biol. 2018, 217, 4025–4048.

(37) van Steensel, B.; Belmont, A. S. Lamina-Associated Domains: Links with Chromosome Architecture, Heterochromatin, and Gene Repression. Cell 2017, 169, 780–791.

(38) Chen, Y.; Belmont, A. S. Genome organization around nuclear speckles. Curr. Opin. Genet. Dev. 2019, 55, 91–99.

(39) Bajpai, G.; Pavlov, D. A.; Lorber, D.; Volk, T.; Safran, S. Mesoscale phase separation of chromatin in the nucleus. Biophysical Journal 2020, 118, 549a.

(40) Laghmach, R.; Di Pierro, M.; Potoyan, D. A. The interplay of chromatin phase separation and lamina interactions in nuclear organisation. bioRxiv 2021,

(41) Chiang, M.; Michieletto, D.; Brackley, C. A.; Rattanavirotkul, N.; Mohammed, H.; Marenduzzo, D.; Chandra, T. Polymer Modeling Predicts Chromosome Reorganization in Senescence. Cell Rep. 2019, 28, 3212–3223.e6.

(42) Maji, A.; Ahmed, J. A.; Roy, S.; Chakrabarti, B.; Mitra, M. K. A Lamin-Associated chromatin model for chromosome organization. Biophysical Journal 2020, 118, 3041–3050.

(43) Li, Q.; Tjong, H.; Li, X.; Gong, K.; Zhou, X. J.; Chiolo, I.; Alber, F. The threedimensional genome organization of Drosophila melanogaster through data integration. Genome Biol. 2017, 18, 1–22.

(44) Paulsen, J.; Sekelja, M.; Oldenburg, A. R.; Barateau, A.; Briand, N.; Delbarre, E.; Shah, A.; Sørensen, A. L.; Vigouroux, C.; Buendia, B.; Collas, P. Chrom3D: three-dimensional genome modeling from Hi-C and nuclear lamin-genome contacts. Genome Biol. 2017, 18, 21.

(45) Dekker, J.; Marti-Renom, M. A.; Mirny, L. A. Exploring the three-dimensional organization of genomes: interpreting chromatin interaction data. Nature Reviews Genetics 2013, 14, 390–403.

(46) Brackey, C. A.; Marenduzzo, D.; Gilbert, N. Mechanistic modeling of chromatin folding to understand function. Nat. Methods 2020, 17, 767–775.

(47) Zhou, R.; Gao, Y. Q. Polymer models for the mechanisms of chromatin 3D folding: review and perspective. Physical Chemistry Chemical Physics 2020, 22, 20189–20201.

(48) Lin, X.; Qi, Y.; Latham, A. P.; Zhang, B. Multiscale modeling of genome organization with maximum entropy optimization. J. Chem. Phys. 2021, 155, 010901.

(49) Bohn, M.; Heermann, D. W. Diffusion-driven looping provides a consistent provides a consistent framework for chromatin organization. PLoS One 2010, 5, e12218.

(50) Barbieri, M.; Chotalia, M.; Fraser, J.; Lavitas, L.-M.; Dostie, J.; Pombo, A.; Nicodemi, M. Complexity of chromatin folding is captured by the strings and binders switch model. Proc. Natl. Acad. Sci. 2012, 109, 16173–16178.

(51) Gürsoy, G.; Xu, Y.; Kenter, A. L.; Liang, J. Computational construction of 3D chromatin ensembles and prediction of functional interactions of alpha-globin locus from 5C data. Nucleic Acids Res. 2017, 45, 11547–11558.

(52) Di Pierro, M.; Cheng, R. R.; Lieberman Aiden, E.; Wolynes, P. G.; Onuchic, J. N. De novo prediction of human chromosome structures: Epigenetic marking patterns encode genome architecture. Proc. Natl. Acad. Sci. 2017, 114, 12126–12131.

(53) Erdel, F.; Rippe, K. Formation of Chromatin Subcompartments by Phase Separation. Biophys. J. 2018, 114, 2262–2270.

(54) MacPherson, Q.; Beltran, B.; Spakowitz, A. J. Bottom–up modeling of chromatin segregation due to epigenetic modifications. Proc. Natl. Acad. Sci. 2018, 115, 12739 LP –12744.

(55) Nuebler, J.; Fudenberg, G.; Imakaev, M.; Abdennur, N.; Mirny, L. A. Chromatin organization by an interplay of loop extrusion and compartmental segregation. Proc. Natl. Acad. Sci. 2018, 115, E6697–E6706.

(56) Qi, Y.; Zhang, B. Predicting three-dimensional genome organization with chromatin states. PLOS Comput. Biol. 2019, 15, e1007024.

(57) Huang, K.; Li, Y.; Shim, A. R.; Virk, R. K.; Agrawal, V.; Eshein, A.; Nap, R. J.; Almassalha, L. M.; Backman, V.; Szleifer, I. Physical and data structure of 3D genome. Science advances 2020, 6, eaay4055.

(58) Laghmach, R.; Di Pierro, M.; Potoyan, D. A. Mesoscale Liquid Model of Chromatin Recapitulates Nuclear Order of Eukaryotes. Biophys. J. 2020, 118, 2130–2140.

(59) Zhang, B.; Wolynes, P. G. Genomic Energy Landscapes. Biophys. J. 2017, 112, 427–433.

(60) Zhang, B.; Wolynes, P. G. P. Topology, structures, and energy landscapes of human chromosomes. Proc. Natl. Acad. Sci. 2015, 112, 6062–6067.

(61) Latham, A. P.; Zhang, B. Improving Coarse-Grained Protein Force Fields with Small-Angle X-ray Scattering Data. J. Phys. Chem. B 2019, 123, 1026–1034.

(62) Xie, W. J.; Zhang, B. Learning the Formation Mechanism of Domain-Level Chromatin States with Epigenomics Data. Biophys. J. 2019, 116, 2047–2056.

(63) Lafontaine, D. L. J.; Riback, J. A.; Bascetin, R.; Brangwynne, C. P. The nucleolus as a multiphase liquid condensate. Nat. Rev. Mol. Cell Biol. 2021, 22, 165–182.

(64) Wang, Y.; Zhang, Y.; Zhang, R.; van Schaik, T.; Zhang, L.; Sasaki, T.; Peric-Hupkes, D.; Chen, Y.; Gilbert, D. M.; van Steensel, B.; Belmont, A. S.; Ma, J. SPIN reveals genome-wide landscape of nuclear compartmentalization. Genome Biol. 2021, 22, 36.

(65) Qi, Y.; Zhang, B. Chromatin Network Retards Droplet Coalescence. bioRxiv 2021,

(66) Safran, S.; Rehovot, I. Statistical Thermodynamics Of Surfaces, Interfaces And Membranes; Frontiers in physics; Avalon Publishing, 1994.

(67) Brackley, C. A.; Liebchen, B.; Michieletto, D.; Mouvet, F.; Cook, P. R.; Marenduzzo, D. Ephemeral Protein Binding to DNA Shapes Stable Nuclear Bodies and Chromatin Domains. Biophys. J. 2017, 112, 1085–1093.

(68) Söding, J.; Zwicker, D.; Sohrabi-Jahromi, S.; Boehning, M.; Kirschbaum, J. Mechanisms for Active Regulation of Biomolecular Condensates. Trends Cell Biol. 2020, 30, 4–14.

(69) Krietenstein, N.; Abraham, S.; Venev, S. V.; Abdennur, N.; Gibcus, J.; Hsieh, T.-h. S.; Parsi, K. M.; Yang, L.; Maehr, R.; Mirny, L. A.; Dekker, J.; Rando, O. J. Ultrastructural Details of Mammalian Chromosome Architecture. Mol. Cell 2020, 78, 554–565.e7.

(70) Xiong, K.; Ma, J. Revealing Hi-C subcompartments by imputing inter-chromosomal chromatin interactions. Nat. Commun. 2019, 10, 1–12.

(71) Zane, L.; Chapus, F.; Pegoraro, G.; Misteli, T. HiHiMap: single-cell quantitation of histones and histone posttranslational modifications across the cell cycle by high-throughput imaging. Mol. Biol. Cell 2017, 28, 2290–2302.

(72) Boyle, S.; Gilchrist, S.; Bridger, J. M.; Mahy, N. L.; Ellis, J. A.; Bickmore, W. A. The spatial organization of human chromosomes within the nuclei of normal and emerin-mutant cells. Human molecular genetics 2001, 10, 211–9.

(73) Farley, K. I.; Surovtseva, Y.; Merkel, J.; Baserga, S. J. Determinants of mammalian nucleolar architecture. Chromosoma 2015, 124, 323–331.

(74) Spector, D. L.; Lamond, A. I. Nuclear speckles. Cold Spring Harb. Perspect. Biol. 2011, 3, 1–12.

(75) Shin, Y.; Chang, Y. C.; Lee, D. S.; Berry, J.; Sanders, D. W.; Ronceray, P.; Wingreen, N. S.; Haataja, M.; Brangwynne, C. P. Liquid Nuclear Condensates Mechanically Sense and Restructure the Genome. Cell 2018, 175, 1481–1491.e13.

(76) Kim, J.; Han, K. Y.; Khanna, N.; Ha, T.; Belmont, A. S. Nuclear speckle fusion via longrange directional motion regulates speckle morphology after transcriptional inhibition. J. Cell Sci. 2019, 132, jcs226563.

(77) Eils, R.; Gerlich, D.; Tvarusko, W.; Spector, D. L.; Misteli, T. Quantitative imaging of pre-mRNA splicing factors in living cells. Mol. Biol. Cell 2000, 11, 413–418.

(78) Khanna, N.; Zhang, Y.; Lucas, J. S.; Dudko, O. K.; Murre, C. Chromosome dynamics near the sol-gel phase transition dictate the timing of remote genomic interactions. Nat. Commun. 2019, 10, 1–13.

(79) Eshghi, I.; Eaton, J. A.; Zidovska, A. Interphase Chromatin Undergoes a Local Sol-Gel Transition upon Cell Differentiation. Phys. Rev. Lett. 2021, 126, 228101.

(80) Walter, J.; Schermelleh, L.; Cremer, M.; Tashiro, S.; Cremer, T. Chromosome order in HeLa cells changes during mitosis and early G1, but is stably maintained during subsequent interphase stages. J. Cell Biol. 2003, 160, 685–697.

(81) Bronstein, I.; Israel, Y.; Kepten, E.; Mai, S.; Shav-Tal, Y.; Barkai, E.; Garini, Y. Transient anomalous diffusion of telomeres in the nucleus of mammalian cells. Phys. Rev. Lett. 2009, 103, 1–4.

(82) Chubb, J. R.; Boyle, S.; Perry, P.; Bickmore, W. A. Chromatin motion is constrained by association with nuclear compartments in human cells. Curr. Biol. 2002, 12, 439–445.

(83) Bronshtein, I.; Kepten, E.; Kanter, I.; Berezin, S.; Lindner, M.; Redwood, A. B.; Mai, S.; Gonzalo, S.; Foisner, R.; Shav-Tal, Y.; Garini, Y. Loss of lamin A function increases chromatin dynamics in the nuclear interior. Nat. Commun. 2015, 6, 8044.

(84) Strom, A. R. et al. HP1α is a chromatin crosslinker that controls nuclear and mitotic chromosome mechanics. Elife 2021, 10.

(85) Boninsegna, L.; Yildirim, A.; Polles, G.; Quinodoz, S. A.; Finn, E. Integrative Genome Modeling Platform reveals essentiality of rare contact events in 3D genome organizations. 2021, 1–50.

(86) Boettiger, A. N.; Bintu, B.; Moffitt, J. R.; Wang, S.; Beliveau, B. J.; Fudenberg, G.; Imakaev, M.; Mirny, L. A.; Wu, C. T.; Zhuang, X. Super-resolution imaging reveals distinct chromatin folding for different epigenetic states. Nature 2016, 529, 418–422.

(87) Handwerger, K. E.; Cordero, J. A.; Gall, J. G. Cajal bodies, nucleoli, and speckles in the Xenopus oocyte nucleus have a low-density, sponge-like structure. Molecular biology of the cell 2005, 16, 202–211.

(88) Lee, H. H.; Kim, H. S.; Kang, J. Y.; Lee, B. I.; Ha, J. Y.; Yoon, H. J.; Lim, S. O.; Jung, G.; Suh, S. W. Crystal structure of human nucleophosmin-core reveals plasticity of the pentamer–pentamer interface. Proteins: Structure, Function, and Bioinformatics 2007, 69, 672–678.

(89) Scherl, A.; Couté, Y.; Déon, C.; Callé, A.; Kindbeiter, K.; Sanchez, J.-C.; Greco, A.; Hochstrasser, D.; Diaz, J.-J. Functional proteomic analysis of human nucleolus. Molecular biology of the cell 2002, 13, 4100–4109.

(90) Plimpton, S.; National, L. S. Fast Parallel Algorithms for Short–Range Molecular Dynamics. J. Comput. Phys. 1995, 117, 1–42.

(91) Durand, N. C.; Shamim, M. S.; Machol, I.; Rao, S. P. S. P.; Huntley, M. H.; Lander, E. S.; Aiden, E. L. Juicer Provides a One-Click System for Analyzing Loop-Resolution Hi-C Experiments. Cell Syst. 2016, 3, 95–98.

(92) Ester, M.; Kriegel, H.-P.; Sander, J.; Xu, X., et al. A density-based algorithm for discovering clusters in large spatial databases with noise. Kdd. 1996; pp 226–231.

